## Supplementary Tables for "GCN2 eIF2 kinase promotes prostate cancer by maintaining amino acid homeostasis"

**Supplementary Table 1.** Summary of transcriptome analysis of LNCaP cells treated with GCN2iB.

| Treatment | Time (hours) | Number of Transcripts[a,b] | | | | | |
|---|---|---|---|---|---|---|---|
| | | Increase | ≥ 2X Increase | Decrease | ≥ 2X Increase | Total | ≥ 2X Total |
| GCN2iB | 6 | 2075 | 2 | 1454 | 25 | 3529 | 27 |
| GCN2iB | 24 | 4205 | 253 | 3738 | 391 | 7943 | 644 |

[a]Unique Ensemble transcript IDs
[b]False discovery rate (fdr), Benjamini-Hochberg adjusted *p*-value ≤ 0.05

**Supplementary Table 2**. siRNAs used in PCa growth studies.

| siRNA Name | siRNA Sequence[a] | Vendor | Catalog Number |
|---|---|---|---|
| Non-Targeting Control | UGGUUUACAUGUCGACUAA | Dharmacon | D-001810-10-20 |
| | UGGUUUACAUGUUGUGUGA | Dharmacon | |
| | UGGUUUACAUGUUUUCUGA | Dharmacon | |
| | UGGUUUACAUGUUUUCCUUA | Dharmacon | |
| Non-Targeting Control | UGGUUUACAUGUCGACUAA | Dharmacon | D-001910-10-05 |
| | UGGUUUACAUGUUUUCUGA | Dharmacon | |
| | UGGUUUACAUGUUUUCCUA | Dharmacon | |
| | UCCUUUACAUGUUGUGUGA | Dharmacon | |
| HRI-1 | GCACAAACUUCACGUUACU | Dharmacon | J-005007-05 |
| HRI-2 | GAUUAAGGGUGCAACUAAA | Dharmacon | J-005007-06 |
| HRI-3 | GCAGAAAUCCAGGUGUUAA | Dharmacon | J-005007-07 |
| HRI-4 | GGUCAGGAAUAAAUUAGAU | Dharmacon | J-005007-08 |
| PKR-1 | GUAAGGGAACUUUGCGAUA | Dharmacon | J-003527-09 |
| PKR-2 | GCGAGAAACUAGACAAAGU | Dharmacon | J-003527-10 |
| PKR-3 | CGACCUAACACAUCUGAAA | Dharmacon | J-003527-11 |
| PKR-4 | CCACAUGAUAGGAGGUUUA | Dharmacon | J-003527-12 |
| PERK-1 | CCAAUGGGAUAGUGACGAA | Dharmacon | J-004883-09 |
| PERK-2 | GGUAGGAUCUGAUGAAUUU | Dharmacon | J-004883-10 |
| PERK-3 | GCAAUUAGCCUUAAGUUGU | Dharmacon | J-004883-11 |
| PERK-4 | AAAUUUGGCUGAAAGAUGA | Dharmacon | J-004883-12 |
| GCN2-1 | CAGCAGAAAUCAUGUACGA | Dharmacon | J-005314-05 |
| GCN2-2 | GCAAUUCUGUGGUGCAUAA | Dharmacon | J-005314-06 |
| GCN2-3 | GACCAUCCCUAGUGACUUA | Dharmacon | J-005314-07 |
| GCN2-4 | GGAAAUUGCUAGUUUGUCA | Dharmacon | J-005314-08 |
| GCN2-5 | CCCUUGUCUCGGAUAAAGA | Dharmacon | A-005314-17 |
| GCN2-6 | CUGGAAUAAUGGAAUGUUG | Dharmacon | A-005314-18 |
| GCN2-7 | UCAUUGAGUUUGAAGAAUU | Dharmacon | A-005314-19 |
| GCN2-8 | UCUUUGUCUUCUAAUAGUC | Dharmacon | A-005314-20 |
| ATF4-1 | CAGAUUGGAUGUUGGAGAA | Dharmacon | J-005125-10 |
| ATF4-2 | CGACUUGGAUGCCCUGUUG | Dharmacon | J-005125-11 |
| ATF4-3 | GAAGAACGAGGCUCUAAAA | Dharmacon | J-005125-12 |
| ATF4-4 | GAGUAGGAAGCCAGACUA | Dharmacon | J-005125-13 |
| SLC3A2-1 | GGACCUUACUCCCAACUAC | Dharmacon | J-003542-09 |
| SLC3A2-2 | GAAUGAGCGUUUUCUGGUA | Dharmacon | J-003542-10 |
| SLC3A2-3 | GUUCCAGGUUCGGGACAUA | Dharmacon | J-003542-11 |
| SLC3A2-4 | GCGCAGAAGUGGUGGCACA | Dharmacon | J-003542-12 |

[a]siRNAs used throughout these studies are highlighted in red.

**Supplementary Table 3.** Oligonucleotides used for library prep, PCR, and qRT-PCR analysis.

| Primer Name | Primer Sequence |
|---|---|
| 4F2-FP1 | 5’-TCTGGACCTTACTCCCAACTA-3’ |
| 4F2-RP1 | 5’-ATCAAGAGCCTGTCTTCACTG-3’ |
| HsATF4-FP | 5’-TCAAACCTCATGGGTTCTCC-3’ |
| HsATF4-RP | 5’-GTGTCATCCAACGTGGTCAG-3’ |
| sgRNA_F | 5’-GCTTTATATATCTTGTGGAAAGGACGAAACACC-3’ |
| sgRNA_R | 5’-CAAGTTGATAACGGACTAGCCTT-3’ |
| IlluminaAd_F | 5’-ACACTCTTTCCCTACACGACGCTCTTCCGATCTTGCTTTATATATCTTGTGGA-3’ |
| IlluminaAd_F | 5’- GACTGGAGTTCAGACGTGTGCTCTTCCGATCTGCCAAGTTGATAACGGACTAGCCTT-3’ |
| 5’-adaptor | 5’-rGrUrUrCrArGrArGrUrUrCrUrArCrArGrUrCrCrGrArCrGrArUrC-3’ |
| RT primer | 5’-AGACGTGTGCTCTTCCGATCT-3’ |
| Illumina Multiplex Primer | 5’-AATGATACGGCGACCACCGAGATCTACACGTTCAGAGTTCTACAGTCCGA-3’ |
| Illumina Barcode Primer | 5’-CAAGCAGAAGACGGCATACGAGATGCCTAAGTGACTGGAGTTCAGACGTGTGCTCTTCCGATCT-3’ |
| tRNAHis-FW | 5’-GCCGTGATCGTATAGTGG-3’ |
| tRNAHis-REV | 5’-GAGAATTCCATGGTGCCGTGACTCGG-3’ |

**Supplementary Table 4.** Composition and dietary intake of essential amino acids from the drinking water.

| Amino Acids | Molecular Mass (g/mol) | Percentage (%) | Dietary intake (mg/g/day) |
|---|---|---|---|
| L-Arginine hydrochloride | 211 | 22.6% | 0.340 |
| L-Cystine | 240 | 4.3% | 0.064 |
| L-Histidine hydrochloride | 210 | 7.5% | 0.113 |
| L-Isoleucine | 131 | 9.4% | 0.141 |
| L-Leucine | 131 | 9.4% | 0.141 |
| L-Lysine hydrochloride | 183 | 13.0% | 0.195 |
| L-Methionine | 149 | 2.7% | 0.041 |
| L-Phenylalanine | 165 | 5.9% | 0.089 |
| L-Threonine | 119 | 8.5% | 0.128 |
| L-Tryptophan | 204 | 1.8% | 0.027 |
| L-Tyrosine | 181 | 6.4% | 0.097 |
| L-Valine | 117 | 8.4% | 0.126 |
